# Cerulean Protein Tagging Enhances Immunogenicity and Osimertinib Response of Transplantable Murine EGFR Mutant Lung Cancer Cell Lines

**DOI:** 10.64898/2026.11.17.767321

**Authors:** Jasalyn T Avila, Trista K Hinz, Andre C Navarro, Kari J Kurppa, Raphael A Nemenoff, Lynn E Heasley

## Abstract

Lung cancers bearing mutations in epidermal growth factor receptors (EGFR) exhibit marked, but incomplete responses to tyrosine kinase inhibitors (TKIs) like osimertinib. We previously developed genetically-engineered mouse models (GEMMs) for EGFR exon 19 deletion (del19) and L860R (mouse equivalent of L858R). Sequences encoding mC3 cerulean fluorescent protein were included to mark emergent tumor cells. Orthotopic implantation of mEGFRdel19 cell lines into syngeneic C57BL/6J mice yielded tumors that were eliminated upon osimertinib therapy which is not typically observed in patients while mEGFR-L860R cell lines failed to establish tumors. Herein, mEGFRdel19 and mEGFR-L860R cell lines formed tumors in non-induced mEGFRdel19 and mEGFR-L860R GEMMs and green fluorescent protein-transgenic mice, supporting mC3-derived epitopes as mediators of immunoelimination. CRISPR-Cas9 mediated disruption of the mC3 cerulean sequence in established mEGFR cell lines yielded two independent mC3-deleted cell lines that formed orthotopic tumors and exhibited partial osimertinib responses. These studies provide a cautionary tale regarding use of fluorescent proteins in murine cancer models and reinforce host immunity as a participant in anti-tumor TKI responses initiated. These mC3-deleted cell lines serve as improved models to explore mechanisms of incomplete TKI response, acquired resistance and cancer cell interaction with the host immune microenvironment.

## Introduction

The identification of lung cancer subsets driven by oncogenic receptor tyrosine kinases (RTKs) including mutated EGFR and rearranged ALK, ROS1 and RET genes coupled with the development of highly specific tyrosine kinase inhibitors (TKIs) provides a personalized medicine approach to this disease. Therapeutic responses to TKIs are impressive, but these agents generally induce partial tumor shrinkage where the residual disease serves as a reservoir for selection and outgrowth of TKI-resistant cancer cells that ultimately yield treatment failure [1, 2]. Moreover, RTK-driven lung cancers fail to exhibit responsiveness to anti-PD1/PD-L1 axis inhibitors as single agents and combination with TKIs yields significant toxicity [3, 4]. Despite the lack of efficacy of existing immune therapies in RTK-driven lung cancers, studies demonstrate the active participation of adaptive immunity in the response of EGFR, ALK and KRAS-driven lung cancers to oncogene-targeted agents [5–7]. A mechanistic understanding of how adaptive immune cells are instructed to positively contribute to the TKI response remains ill-defined and clarification may open novel avenues to enhance the durability of precision oncology therapeutic response.

To explore the mechanisms by which adaptive immune cells, especially CD8+ T cells, are recruited and contribute to the anti-tumor TKI responses, we developed genetically engineered mouse models (GEMMs) as tools to study EGFR, ALK and RET-driven lung cancers [6–9]. We used these novel GEMMs to generate stable tumor-derived cell lines that can be orthotopically transplanted into lungs of syngeneic C57BL/6 murine hosts. Regarding oncogenic EGFR, murine lung cancer cell lines bearing an EGFR exon 19 deletion (mEGFRdel19) or the murine equivalent of the EGFR-L858 mutation (mEGFR-L860R) were established. Orthotopic tumors established in syngeneic hosts from mEGFRdel19 lines exhibited marked shrinkage with osimertinib treatment. In fact, complete tumor elimination was observed. This is certainly not a typical result observed in human patients where partial tumor shrinkage and frequent acquisition of acquired resistance predominate. The complete response observed with the mEGFRdel19 cell lines reduces their utility for assessing mechanisms mediating on-treatment acquired resistance. Moreover, the mEGFR-L860R cell lines failed to establish orthotopic tumors in syngeneic hosts despite induction of primary lung tumors in the GEMM following adeno-Cre administration.

The murine EGFRdel19 and -L860R sequences in the previously reported GEMMs [7] were engineered with a 3’ T2A (*Thosea asigna* virus 2A) peptide-encoding sequence [10] followed by mCerulean3 (mC3) sequences to permit tagging of the emergent lung tumor cells. The mCerulean family of fluorescent proteins are derived from serial amino acid substitutions of the *Aequorea victoria* green fluorescent protein (GFP) [11]. While GFP and GFP-expressing murine cancer lines have been shown to be antigenic in C57BL/6 hosts [12], there are few reports on mCerulean proteins. However, the antigenic epitope within GFP that is presented on HLA proteins in C57BL/6 mice has been identified [13] and is predicted to be conserved in the mC3 protein. Herein, we tested the hypothesis that the failure of mEGFR-L860R cell lines to form tumors in syngeneic hosts and the complete response of mEGFRdel19 tumors to the 3^rd^ generation TKI, osimertinib, is dictated by mC3 functioning as a neoepitope in C57BL/6 hosts. In support, both mEGFR-L860R and mEGFRdel19 cell lines form orthotopic tumors in GFP-transgenic C57BL/6 mice with evidence of stable residual disease and progression following continuous osimertinib treatment. Moreover, parental mEGFR cell lines in which mC3 was deleted by CRISPR/Cas9 editing formed viable orthotopic tumors in syngeneic hosts and exhibited residual disease and frequent progression with osimertinib therapy. While this study serves as a cautionary tale regarding the use of fluorescent proteins to tag cancer cells, it also reinforces the contribution of adaptive immunity to TKI responsiveness. Finally, these re-engineered mEGFRdel19 and mEGFR-L860R cell lines will be valuable tools for addressing mechanisms of residual disease and acquired resistance in murine preclinical models.

## Results

We previously reported genetically engineered mouse models (GEMMs) that permit Cre-inducible expression of murine EGFRdel19 and -L860R (murine equivalent of human L858R) [7]. In Figure 1, bronchial instillation of Adeno-Cre yields multiple lung tumors in both EGFRdel19 and EGFR-L860R models that undergo strong shrinkage responses with osimertinib therapy (5 days on and 2 days off). After 49 days of osimertinib treatment, μCT revealed evidence of on-treatment progression of some tumor foci in the EGFRdel19 mouse (upper panel). Also, terminating osimertinib treatment in the EGFR-L860R mouse after ∼2 months of treatment resulted in marked regrowth of lung tumors detected by μCT (Fig. 1, lower panel). Thus, EGFR mutant lung tumors induced by Adeno-Cre instillation in the GEMMs exhibit partial responses and residual disease, key features that limit the long-term durability of TKI therapy in patients.

**Figure 1.**
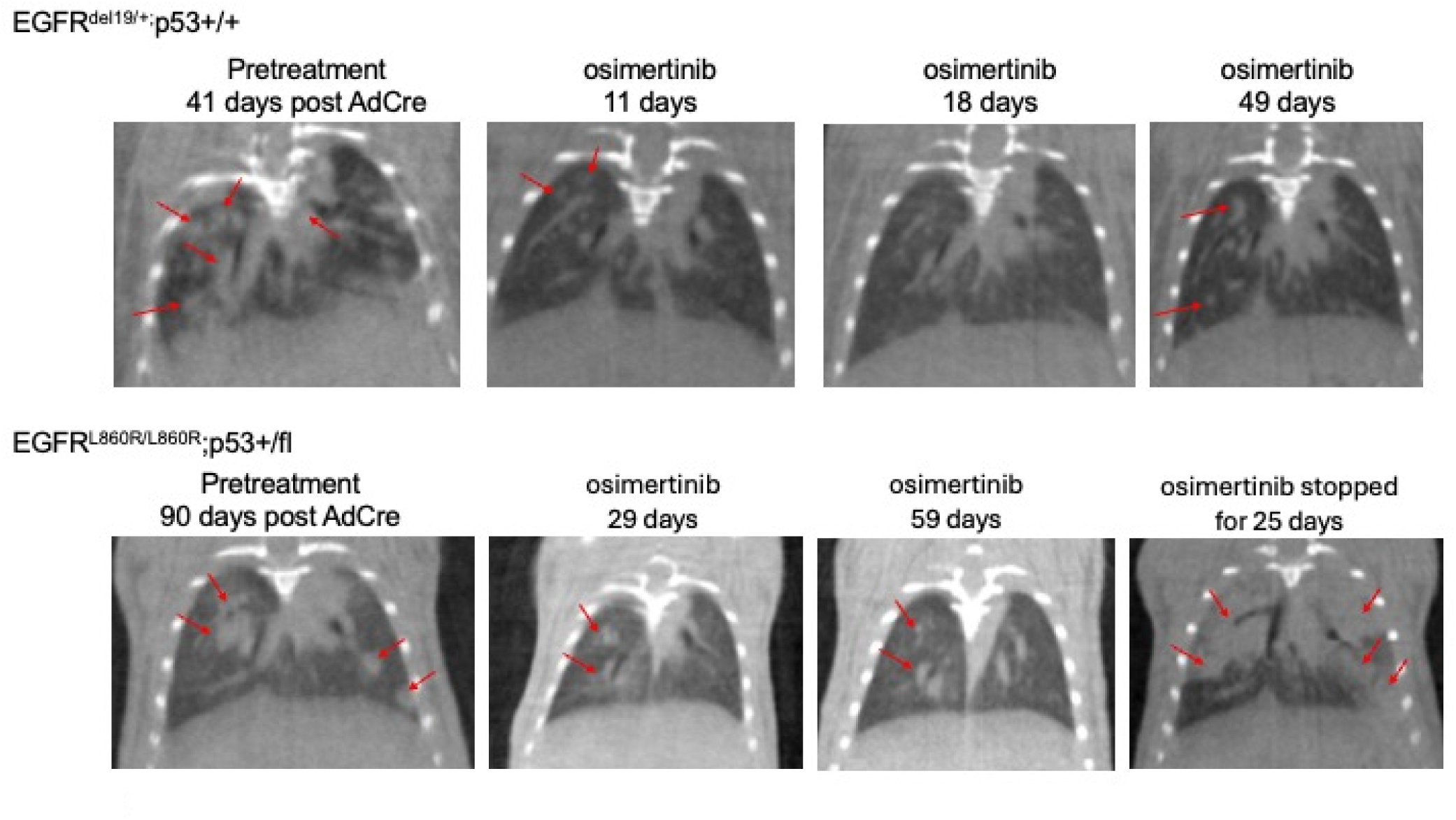
Osimertinib response in primary lung tumors induced in mEGFRdel19 and mEGFR-L860R GEMMs. The previously reported EGFR GEMM models were intra-tracheally instilled with Adeno-Cre virus as described [7]. When lung tumors were evident as assessed by μCT (41 days for the mEGFRdel19 mouse, 90 days for the mEGFR-L860R mouse), osimertinib therapy (5 mg/kg for 5 days then 2 days off) was initiated. The figure shows a pre-treatment scan for each mouse and serial scans after the indicated number of days of osimertinib therapy. The representative mEGFRdel19 mouse presented with multiple lung nodules (noted with red arrows) that underwent marked shrinkage with osimertinib treatment and μCT evidence for on-treatment progression of two nodules after 49 days of treatment (see red arrows). The lung tumors in the mEGFR-L860R mouse exhibited a partial response to osimertinib and extensive regrowth after cessation of therapy at day 59.

In our previous report [7], primary Cre-induced lung tumors were used to derive two distinct mEGFRdel19 cell lines, mEGFRdel19.1 and mEGFRdel19.2. Orthotopic implantation of these cell lines into the left lung of syngeneic C57BL/6J mice yielded tumors that strongly responded to administered osimertinib. In fact, osimertinib treatment completely eliminated the tumors as evidenced by failure of tumor regrowth following cessation of TKI treatment [7]. By contrast, a partial osimertinib response was observed when these cell lines were used to establish orthotopic tumors in immune-deficient mice, supporting our premise that adaptive immunity contributes to the TKI response. Also, a cell line (mEGFR-L860R.1) was derived from an L860R primary tumor, but it reproducibly failed to establish orthotopic tumors upon transplantation into syngeneic C57BL/6J hosts [7]. Since publication of this study, we generated additional L860R cell lines (mEGFR-L860R.2, mEGFR-L860R.3 and multiple subclones). Like mEGFR-L860R.1 cells, these new lines failed to successfully establish viable orthotopic lung tumors that increased in size over time in syngeneic hosts (Fig. 2A) but yielded progressing tumors upon inoculation into immune deficient Rag1 hosts (Fig. 2B). This study suggested that adaptive immunity is responsible for the failure of the mEGFR-L860R cell lines to form tumors in syngeneic hosts as well as the complete elimination of mEGFRdel19-derived tumors in response to osimertinib.

**Figure 2.**
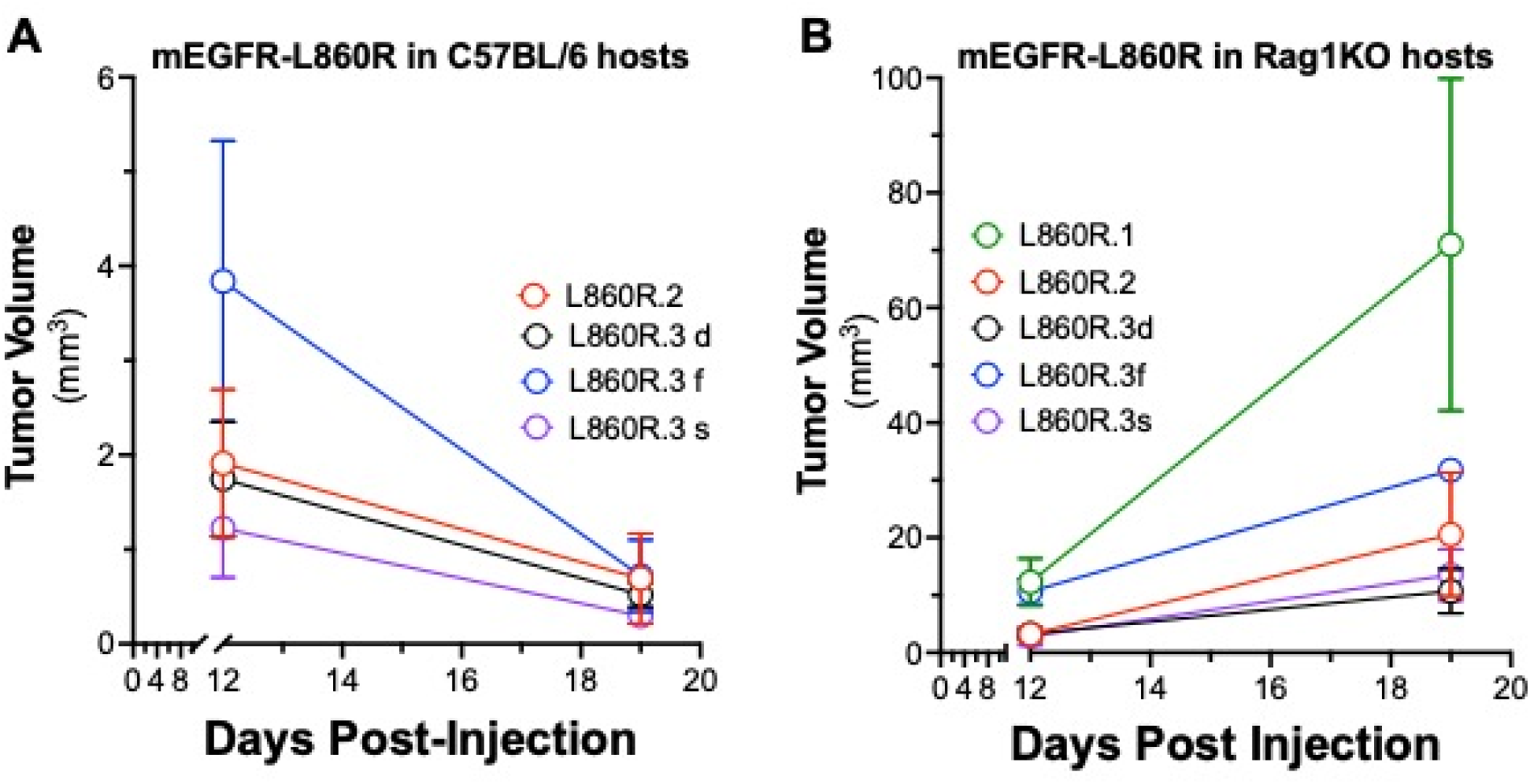
Evidence for spontaneous elimination of mEGFR-L860R lung cancer cell lines in immune-competent murine hosts. The indicated mEGFR-L860R cell lines were inoculated into the left lungs of syngeneic C57BL/6J mice (**A**) or immune-deficient Rag1KO mice (**B**). After twelve and nineteen days, the mice were submitted to μCT imaging to obtain tumor volume measurements. The data are means and SEM of 3 to 5 mice per cell line.

### Murine EGFRdel19 and EGFR-L860R cell lines form orthotopic tumors in non-Cre-induced EGFR GEMMs

Primary mEGFRdel19 and mEGFR-L860R lung tumors induced in the GEMMs exhibit marked osimertinib responses (Fig. 1), yet the tumors are not eliminated and occasionally progress on continuous TKI treatment. Thus, the observation of complete tumor elimination of transplanted orthotopic mEGFRdel19 tumors and the failure of mEGFR-L860R cell lines to establish lung tumors in syngeneic C57BL/6J hosts is unexpected. SNP analysis (Transnetyx, Cordova, TN) of the mEGFRdel19 and mEGFR-L860R cell lines confirmed that they are of C57BL/6 origin (>99%, data not shown). The ability of the mEGFRdel19 and L860R cell lines to form orthotopic tumors in the GEMMs from which they were derived was tested. mEGFRdel19.2 and mEGFR-L860R.1 cells were inoculated into the left lungs of the respective GEMMs that had not been treated with Adeno-Cre and therefore, should phenotypically reflect wild-type mice. Compared to the results with C57BL/6J mice, all injected GEMMs exhibited evidence for tumor establishment by μCT (Fig. 3A and B). Of three mEGFRdel19 GEMM mice bearing mEGFRdel19.2 tumors, all exhibited evidence by μCT of residual disease following ∼10 weeks of osimertinib therapy (Fig. 3A). Moreover, of the mEGFR-L860R.1-derived tumors propagated in EGFR-L860R (open symbols) or EGFRdel19 (closed symbols) GEMMs, 3 exhibited complete osimertinib responses and 2 partial responses (Fig. 3B). The tumor in mouse #122 underwent progression after ∼1 month of treatment while the tumor in mouse #114 which had been undetectable by μCT progressed after 2 months of osimertinib treatment.

**Figure 3.**
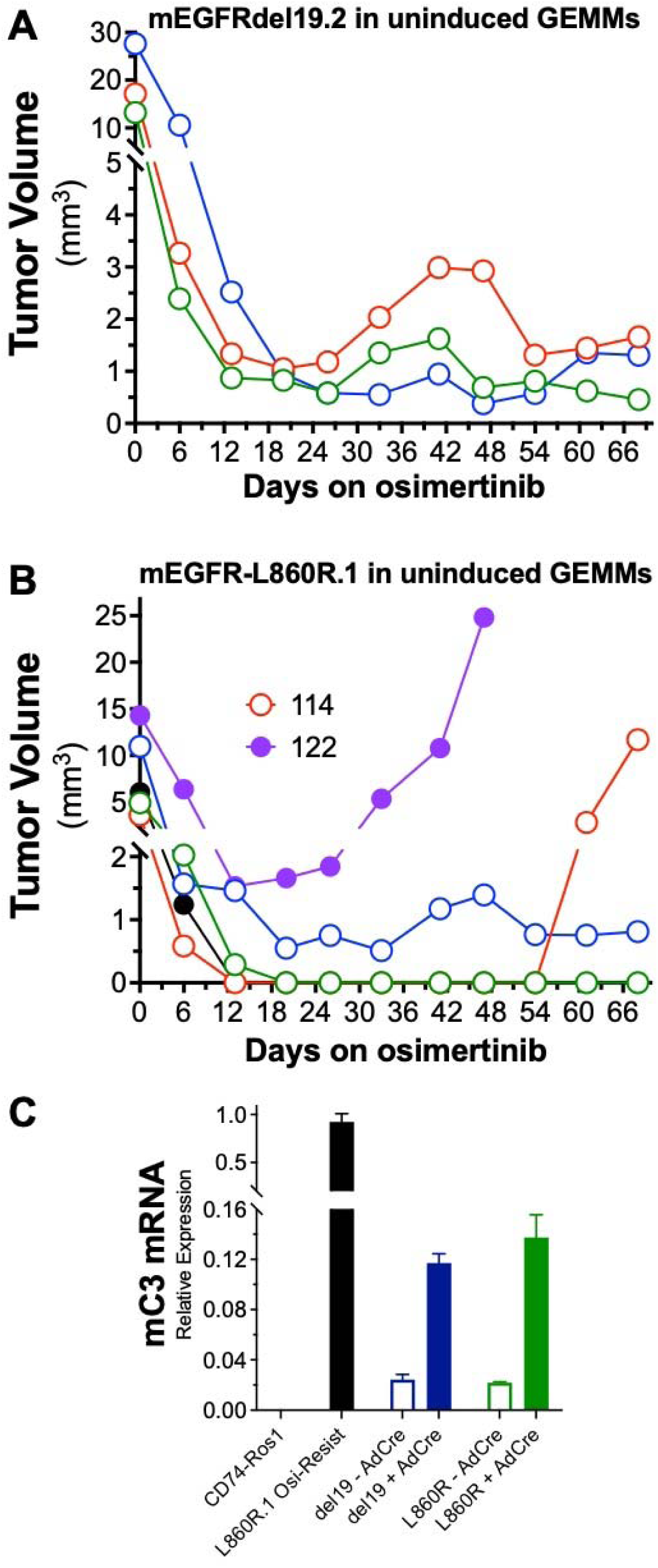
Growth and osimertinib responsiveness of mC3-positive mEGFRdel19.2 and mEGFR-L860R. 1 orthotopic tumors in adeno-Cre naïve mEGFR GEMMs. A, Parental mEGFRdel19.2 and B, mEGFR-L860R.1 cells were orthotopically inoculated into respective adeno-Cre naïve mEGFR GEMMs, and after 10 days, initial tumor volumes were assessed by μCT. The tumor-bearing mice were treated with 5mg/kg osimertinib for 5 days on, 2 days off for the indicated number of days and tumor volumes were monitored by weekly μCT. In B, mouse #122, an mEGFRdel19 GEMM and mouse #114, an mEGFR-L860R GEMM were euthanized on days 43 and 67, respectively and the lung tumors were used to establish osimertinib-resistant cell lines presented in Figure 6. C, Lung fibroblasts were prepared as described in the Materials and Methods from the indicated GEMMs and cultured with or without adeno-Cre. Fibroblasts from a CD74-ROS1 mouse (under development) were used as a negative control and mEGFR-L860R.1 cells were used as a positive control. Total RNA was prepared and mC3 mRNA was measured by quantitative RT-PCR and normalized to GAPDH mRNA levels.

The mEGFR GEMMs were generated with a ROSA-targeting construct in which mCerulean3 (mC3) coding sequences are inserted downstream of the murine EGFR sequences with an intervening T2A-encoding sequence [7]. This results in expression from a single open reading frame of murine EGFRdel19 or EGFR-L860R and mC3 upon excision of the Lox-Neo-Stop-Lox sequence with Cre recombinase. Because pure C57BL/6 mice don’t express mC3, it is possible that this fluorescent protein expressed in the established cell lines is identified as a neoepitope in C57BL/6 hosts. While the GEMMs are not predicted to express mC3 protein until after Adeno-Cre instillation, leaky expression of the mEGFR-mC3 transgene may occur. To test if the mEGFR-mC3 transgene is expressed in cells derived from the GEMM models in the absence of Adeno-Cre treatment, lung fibroblasts were prepared from Adeno-Cre-naïve mEGFRdel19 and -L860R GEMMs as well as from mEGFR-L860R.1 lung cancer cells as a positive control. As a negative control, lung fibroblasts were prepared from a CD74-ROS1 GEMM (under development) in which mC3 sequences were not used in the transgene construct. The mEGFR GEMM-derived fibroblasts were cultured with or without Adeno-Cre particles for 3 days and total RNA was prepared and submitted to quantitative RT-PCR with mC3 primers. The findings in Figure 3C reveal that mC3 RNA is highly expressed in positive control mEGFR-L860R.1 cells and mEGFR GEMM-derived fibroblasts exposed to Adeno-Cre. Importantly, relative to the negative control CD74-ROS1 lung fibroblasts, mC3 mRNA is clearly detectable in EGFR GEMM-derived fibroblasts not incubated with Cre recombinase. Based on this result, mEGFR GEMMs appear to express low levels of mC3 protein prior to Adeno-Cre instillation, likely rendering them immune tolerant to the mC3-positive mEGFRdel19 and -L860R cell lines.

### Propagation of mEGFRdel19 and -L860R cell line-derived tumors in GFP-transgenic mice

The mC3 protein is derived from serial amino acid substitutions of the *Aequorea victoria* green fluorescent protein (GFP) (https://www.fpbase.org/protein/mcerulean3/ and [11]). The amino acid sequence alignment of mC3 and GFP (Fig. 4A) reveals high similarity of the two proteins. In fact, the antigenic epitope within GFP that is presented on HLA proteins in C57BL/6 mice has been identified [13] and is completely conserved in the mC3 protein (Fig. 4A). Thus, mC3-expressing mEGFR cell lines are not predicted to be immunogenic in GFP-transgenic C57BL/6 mice.

**Figure 4.**
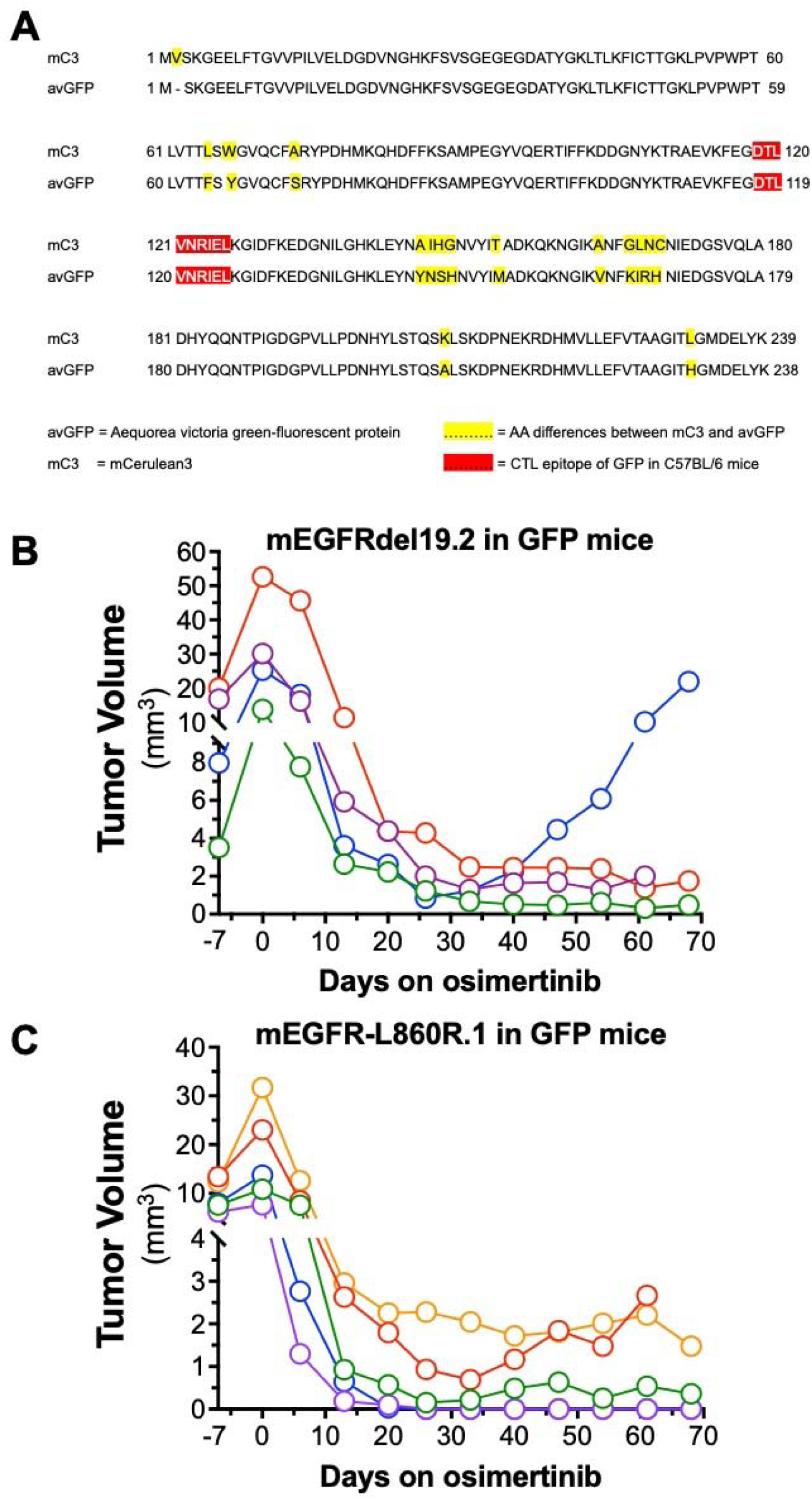
Growth and osimertinib response of mC3-positive mEGFRdel19.2 and mEGFR-L860R.1 tumors propagated in UBC-GFP transgenic mice. **A**, Amino acid sequences for mCerulean 3 (mC3) and Aequorea victoria GFP (avGFP) were aligned with MacVector. The residues highlighted in yellow indicate amino acid differences between mC3 and avGFP. The defined antigenic avGFP epitope [13] in C57BL/6 mice is indicated in red. **B** and **C**, Parental mC3+ mEGFRdel19.2 and mEGFR-L860R.1 cells were orthotopically inoculated into UBC-GFP transgenic mice and after 10 days, initial tumor volumes were measured with μCT. The tumor-bearing mice were treated with 5mg/kg osimertinib for 5 days on, 2 days off for the indicated number of days and tumor volumes were monitored by weekly μCT.

To test this hypothesis, mEGFRdel19.2 and mEGFR-L860R.1 cells were inoculated into the left lungs of GFP-transgenic C57BL/6 mice. Serial μCT imaging of the mice revealed measurable tumors 10 days after orthotopic injection with further increases in tumor size over the next 7 days (Fig. 4B and C). Thus, mC3-positive mEGFR-L860R.1 cells form progressing tumors in GFP-transgenic mice, but not pure C57BL/6 mice (Fig. 2B). The tumor bearing mice were treated with osimertinib and tumor size was monitored by weekly μCT measurements. As shown in Figure 4B and C, all tumors underwent marked shrinkage after 2 weeks of TKI treatment with maximal responses observed within 25 to 30 days. Three of the four mice bearing mEGFRdel19.2 tumors (Fig. 4B) presented with stable residual tumors that were maintained for 61 days of treatment. One tumor (#285, blue symbols) underwent on-treatment progression after 33 days of osimertinib therapy. Of the five mEGFR-L860R.1 injected mice (Fig. 4C), three presented with clear residual tumors after 3 weeks of TKI treatment and two completely responded with no detectable tumor by μCT (Fig. 4C). The mEGFR-L860R.1 tumor in one mouse in red symbols underwent on-treatment progression after 33 days and was euthanized at day 61. These experiments demonstrate that the mC3-positive mEGFRdel19 and -L860R cell lines form tumors in GFP-transgenic C57BL/6 mice that yield residual disease with osimertinib treatment and in several instances, undergo on-treatment progression.

### CRISPR-Cas9-mediated knockout of mC3 in mEGFR cell lines rescues tumor growth and residual disease in C57BL/6J hosts following osimertinib treatment

As an alternative approach to testing the role of mC3 as a neo-epitope in C57BL/6J hosts, mEGFRdel19.2 and mEGFR-L860R.1 cells were submitted to CRISPR-Cas9 editing with a gRNA targeting the mC3 sequence (see Materials and Methods). Cancer cells negative for mC3 were collected by serial flow cytometry gating on the mC3 negative populations as described in the Materials and Methods. The resulting mC3 knockout (mC3-KO) mEGFRdel19.2 and -L860R.1 cell lines were tested for *in vitro* sensitivity to osimertinib relative to the parental lines. As shown in Figure 5A, the mC3-KO cell lines exhibited equivalent osimertinib sensitivity as the parental mC3-positive cells from which they were derived. Both mEGFRdel19.2-mC3KO and mEGFR-L860R.1-mC3KO cell lines formed orthotopic tumors following implantation into lungs of syngeneic C57BL/6J mice and exhibited marked shrinkage following 14 to 21 days of osimertinib therapy (Fig. 5B and C). Of the twelve mice bearing mEGFRdel19.2-mC3KO tumors, all presented with residual, μCT-detectable tumors after 21 days of osimertinib treatment. Moreover, seven of the twelve mEGFRdel19.2-mC3KO tumors underwent clear on-treatment progression (Fig. 5B). This contrasts with the complete elimination of parental mEGFRdel19.2 tumors observed with single-agent osimertinib treatment [7]. Distinct from mC3-positive parental mEGFR-L860R.1 cells (Fig. 2A), the mEGFR-L860R.1-mC3KO cells yielded progressing lung tumors following implantation into C56BL/6J hosts that exhibited strong osimertinib shrinkage responses (Fig. 5C). Of the thirteen tumor-bearing mice, five underwent obvious tumor progression by 2 months of osimertinib treatment. Seven mEGFR-L860R.1-mC3KO tumors yielded detectable residual disease after 60 days of treatment while one tumor showed a complete response that was not detectable by μCT.

**Figure 5.**
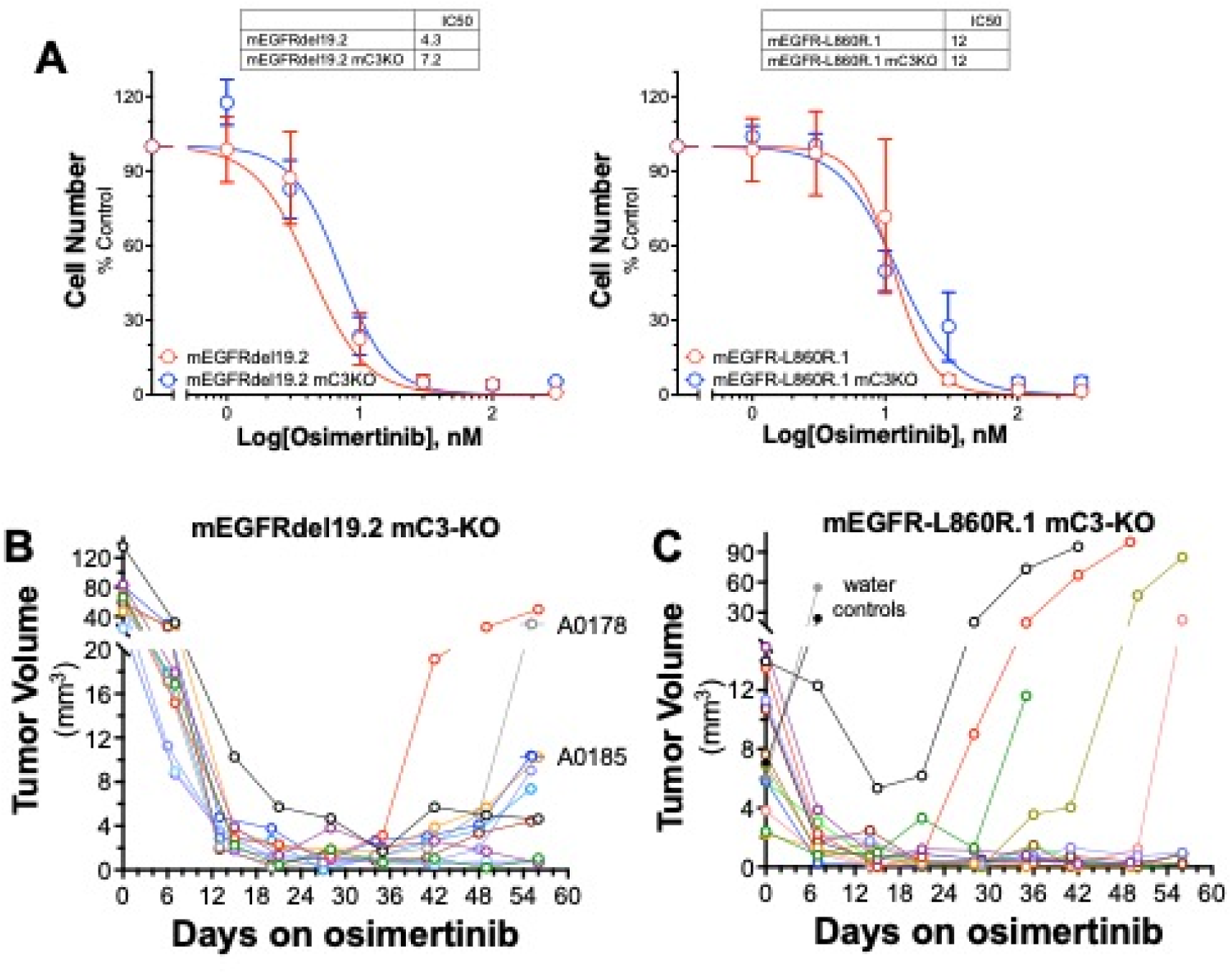
Osimertinib treatment yields residual disease and on-treatment progression in syngeneic hosts bearing orthotopic lung tumors derived from mC3-deficient mEGFRdel19.2 and mEGFR-L860R.1 cells. Parental mEGFRdel19.2 and mEGFR-L860R.1 cells were submitted to CRISPR/Cas9 editing to disrupt mC3 and mC3 knockout cells were collected by cell sorting as described in the Materials and Methods. A, Parental and mC3KO cell lines were seeded in 96-well plates and cell growth was assayed in the presence of increasing osimertinib concentrations. The data are the means and SEM of triplicate determinations, and the IC_50_ values were determined with Prism 10. B, mEGFRdel19.2 mC3-KO and C, mEGFR-L860R.1 mC3-KO cells were orthotopically inoculated into C57BL/6J mice and tumors were allowed to establish for 10 days. After assessing initial tumor volume with μCT (t=0), the mice were treated with osimertinb (5mg/kg) for 5 days on, 2 days off for the indicated number of days and tumor volumes were monitored by weekly μCT. In B, mouse #A0178 and #A0185 were euthanized at day 55 and the tumors progressing on osimertinib were used to establish TKI-resistant cell lines (see Fig. 6). In **C**, two tumor-bearing mice were treated with water instead of osimertinib to demonstrate the rate of tumor growth in mice not treated with TKI.

### Analysis of mEGFR cell lines derived from osimertinib treatment-resistant tumors

The presentation of residual disease and on-treatment progression in mEGFR-mutant lung tumors in GFP transgenic mice and mEGFR-mC3-KO tumors in C57BL/6 mice provide new models for investigating and combatting acquired TKI resistance in an immune-competent setting. Two distinct mEGFRdel19.2-mC3-KO tumors that progressed on osimertinib in C57BL/6 hosts were dissected upon completion of the experiment shown in Figure 5B and used to establish osimertinib-resistant cell lines as described in the Materials and Methods. These TKI-resistant cell lines are referred to as mEGFRdel19.2-mC3-KO-OsiR A0178 and mEGFRdel19.2-mC3-KO-OsiR A0185. Also, two mEGFR-L860R.1 tumors propagated in un-induced mEGFR GEMMs (Fig. 3B) that progressed on continuous osimertinib treatment were similarly collected and used to establish the osimertinib-resistant cell lines, mEGFR-L860R.1-OsiR#114 and mEGFR-L860R.1-OsiR#122. The TKI-resistant cultures were initiated and maintained in medium supplemented with 200 nM osimertinib to retain the resistant state. Compared to parental mEGFRdel19.2-mC3-KO and mEGFR-L860R.1-mC3-KO cells, the osimertinib-resistant cell lines exhibited insensitivity to osimertinib and the distinct EGFR-ERBB2 inhibitor, afatinib (Fig. 6A).

**Figure 6.**
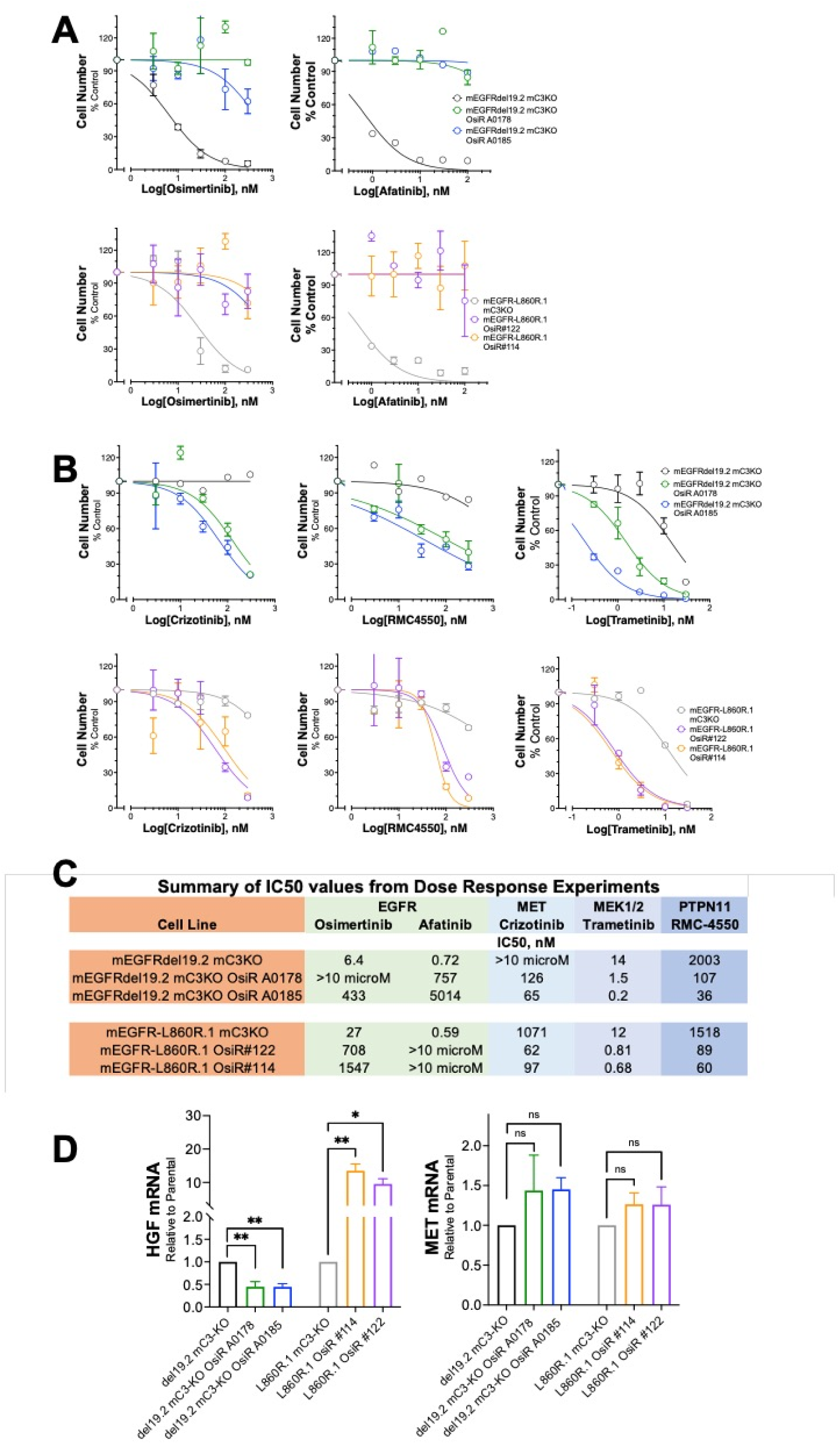
Acquired sensitivity to inhibitors of MET, PTPN11 and MEK in mEGFR cell lines derived from osimertinib-resistant lung tumors. The mEGFRdel19.2 mC3-KO lung tumors in mice labeled A0178 and A0185 (see Fig. 5B) and mEGFR-L860R.1 lung tumors in Adeno-Cre naïve mice #113 and #122 (Fig. 3B) were microdissected, dissociated and cultured in the presence of 200 nM osimertinib until stable cultures were established. In **A** and **B**, the indicated osimertinib resistant (OsiR) cell lines in medium containing 200 nM osimertinib were seeded into 96-well plates with mEGFRdel19.2 mC3KO and mEGFR-L860R.1 mC3KO cells as controls. The indicated drugs were added as 2X concentrates in growth medium and the plates were incubated for 5-7 days. Cell numbers were assayed with CyQUANT reagent as described in the Materials and Methods. The data are the means and SEM of triplicate measurements. **C**, the IC_50_ values from the dose-response curves in **A** and **B** were calculated with Prism 10 and tabulated. **D**, Total RNA was prepared from the control and OsiR cell lines and submitted to quantitative RT-PCR as described in the Materials and Methods and normalized to GAPDH mRNA levels. The data are the means and SEM of three independent assays. Differences between the control and OsiR cell lines were assessed by 1-way ANOVA with correction for multiple comparisons.

We previously reported the generation and analysis of a mEGFRdel19 cell line (mEGFRdel19.1-616) established from an orthotopic tumor that acquired osimertinib resistance when propagated in an immune-deficient nu/nu mouse [7]. In this cell line, osimertinib resistance was driven in part by bypass signaling through an HGF-MET pathway as evidenced by acquired sensitivity to the MET inhibitor, crizotinib. The novel osimertinib-resistant mEGFR cell lines generated herein, but not the parental mC3-KO cell lines, were likewise sensitive to crizotinib (Fig. 6B). Expression of HGF mRNA was increased in mEGFR-L860R.1-OsiR#114 and #122 cells relative to mEGFR-L860R.1-mC3-KO cells (Fig. 6D). By contrast, decreased HGF mRNA was noted in the mEGFRdel19.2-mC3-KO-OsiR A0178 and mEGFRdel19.2-mC3-KO-OsiR A0185 cells relative to mEGFRdel19.2-mC3-KO cells. MET mRNA levels (Fig. 6D) were not altered in any of the osimertinib-resistant cell lines. In addition to acquired crizotinib sensitivity, the osimertinib-resistant cell lines exhibited markedly increased sensitivity to the PTPN11/SHP2 inhibitor, RMC-4550 and the MEK inhibitor, trametinib (Fig. 6B and C). We recently reported [8] a similar sensitization to RMC-4770 and trametinib in murine RET-driven lung cancer cell lines that had acquired *in vivo* resistance to selpercatinib, indicating that increased dependency on PTPN11-MEK-MAPK signaling may be a common event in bypass signaling-mediated TKI resistance.

## Discussion

The present study provides evidence that the cerulean protein, mC3, is causative in the complete elimination of transplanted mEGFRdel19 tumors following osimertinib therapy and the failure of implanted mEGFR-L860R cell lines to successfully form tumors in syngeneic mice. The immunogenicity of fluorescent proteins of the RFP and GFP families has been considered in the literature and, in fact, supported the generation of a GEMM in which defined epitopes for many of these are expressed as a transgene to induce immune tolerance [14]. While there is scant published evidence for an immunogenic role for mCerulean3, studies with closely related mCerulean proteins suggest relatively low antigenicity in C57BL/6 mice, thereby supporting our initial strategy of including mC3 in the EGFR transgene cassette to tag the emergent tumor cells. The findings in Figure 3 that mEGFRdel19.2 and mEGFR-L860R.1 cells can establish tumors in Adeno-Cre naïve GEMMs and exhibit partial responses to osimertinib indicate tolerance to mC3 in these mice. In fact, lung fibroblasts cultured from the GEMMs express readily measurable mC3 mRNA prior to Adeno-Cre exposure, showing that the floxed Neo-stop cassette is leaky. Of note, the defined antigenic epitope [13] in the parental GFP protein from which mCerulean family members are derived is conserved in mC3. Similar to Adeno-Cre naïve GEMMs, mEGFRdel19.2 and mEGFR-L860R.1 cells form viable tumors in UBC-GFP transgenic mice where partial osimertinib responses and on-treatment acquired resistance are observed. Again, this supports immune tolerance to mC3-expressing mEGFR cancer cells in this GFP+ C57BL/6 context. Finally, CRISPR-Cas9-directed disruption of mC3 in the mEGFRdel19.2 and mEGFR-L860R.1 cells rescues the ability of these lines to form viable tumors in C57BL/6 hosts (Fig. 5). Again, both tumor types exhibit partial responses to osimertinib therapy with evidence of on-treatment progression in some mice. These re-engineered mEGFR cell lines are predicted to serve as significantly improved models for addressing important questions regarding partial TKI responses yielding residual disease as well as *in vivo* acquired resistance mechanisms. Alternatively, the original mC3+ mEGFR cell lines can be used in commercially available UBC-GFP transgenic mice to address the same questions. Finally, the mC3+ parental lines propagated in C57BL/6 mice might offer models to identify cancer cell autonomous mechanisms and pathways that blunt the adaptive immune response.

It is notable that mEGFRdel19.2 cells establish viable orthotopic tumors in C57BL/6 mice, but not mEGFR-L860R.1 cells since both cell lines express mC3. The injected mEGFRdel19.2 cells may establish an immune microenvironment that contains reduced tumor-reactive CD8+ T cells relative to mEGFR-L860R.1 tumors until TKI treatment is initiated. Thus, supported by published findings showing increased T cell content in mEGFRdel19 tumors after a 4-day osimertinib treatment [7], TKI administration may stimulate influx of CD8+ T cells, some directed against the mC3/GFP epitope, that subsequently direct the complete elimination of the tumor cells. By contrast, mEGFR-L860R.1 cells may establish an immune microenvironment following orthotopic injection that fails to exclude mC3-reactive T cells such that immune elimination occurs prior to TKI administration. Alternatively, mEGFR-L860R.1 cells may present the mC3 antigenic peptide more effectively than mEGFRdel19 cell lines. In support, differences in baseline antigen presentation machinery are noted in our previous publication (Fig. 6B of [7]). The baseline mRNA expression (mean±SEM) of H2-K1 which presents the antigenic GFP peptide [13] is 1071±257, 1512±308 and 3725±561 CPM in mEGFRdel19.1, mEGFRdel19.2 and mEGFR-L860R.1 cells, respectively while baseline H2-D1 expression is 1833±301, 1447±161 and 2050±326, respectively in the same cell lines. The 2.5 – 3.5-fold increase in H2-K1 expression in mEGFR-L860R.1 cells may significantly enhance their recognition by CD8+ T cells when occupied with the GFP/mC3 epitope.

When parental mEGFR cell lines are propagated as orthotopic tumors in Adeno-Cre naïve GEMMs (Fig. 3A and B) and GFP-transgenic mice (Fig. 4B and C) or mC3-KO cell lines are propagated as tumors in syngeneic hosts (Fig. 5B and C), the residual tumor size following osimertinib therapy and the time to treatment relapse varied significantly among individual mice. In this regard, the orthotopic mEGFRdel19 and mEGFR-L860R models present a spectrum of therapeutic responses similar to that observed in EGFR, ALK, RET and ROS1-positive lung cancer patients treated with TKIs. We wonder what may account for this observed variability in the osimertinib therapeutic responses. While inoculation of mEGFR cell lines into the left lung lobes is performed as uniformly as technically possible, subtle variation in the juxtaposition to blood vessels or other anatomical structures may occur. Moreover, we cannot exclude the stochastic nature of molecular events that drive drug tolerance and acquired resistance. It is notable that the HGF-MET pathway identified as an osimertinib resistance mechanism in a mEGFRdel19 tumor that progressed on TKI treatment in an immune deficient mouse [7] is also identified in multiple mEGFRdel19 and mEGFR-L860R tumor-derived cell lines propagated and treated in immune competent hosts (Fig. 6). Relative to the time to osimertinib treatment failure observed herein (∼1 month or greater), we previously reported on-treatment progression of orthotopic tumors within 2 weeks when orthotopic mEGFRdel19 tumors were propagated in immune-deficient hosts [7]. It is possible that the depth of TKI response and the time to osimertinib progression in the models developed herein are related more to the initial strength and durability of the anti-tumor immune response rather than simple Darwinian selection for resistance-mediating molecular events.

The present findings reinforce the concept set forth in prior reports [6, 7] that the anti-tumor response of TKIs is a composite of direct antagonistic activity on the oncogenic RTK and induction of secondary signaling pathways that recruit adaptive immune cells to the tumor. Identifying ways to enhance adaptive immune contributions during TKI therapy is predicted to increase depth and duration of response. Osimertinib treatment of orthotopic mC3+ mEGFRdel19 tumors propagated in syngeneic mice yields complete elimination [7], presumably due to the concerted action of the TKI and host adaptive immunity directed against mC3. While mC3 antigenicity represents an artificial approach to enhancing immune recognition in tumors treated with oncogene-targeted agents, combining TKIs with agents that enhance host immune recognition may represent an approach to yield deeper and more durable tumor shrinkage responses, perhaps even cures. To date, combining TKIs with anti-PD1/PD-L1 axis inhibitors in clinical studies of RTK-driven lung cancers has not been successful due to significant toxicity [3, 4]. An alternative strategy that is in early stages of clinical evaluation involves deployment of EGFR-targeting CAR T cells as a means to reduce or eliminate drug tolerant persister cells and residual disease. Small early-phase clinical trials with EGFR-targeting CAR T cells are ongoing [15, 16]. This idea is being further pursued pre-clinically where EGFR-targeting CAR NK cells demonstrate activity against EGFR-mutant tumors, especially when combined with TKI [17]. In other pre-clinical studies, CAR T cells targeting TROP2 which is induced in TKI-treated patient tumors yielded significantly prolonged duration of response with evidence of cures [18]. Finally, various approaches involving delivery of vaccines to mutant EGFR epitopes in concert with TKIs have been initiated [19]. Besides anti-tumor vaccines targeting the oncogene, another approach submits surgically resected melanoma and normal specimens to DNA sequencing to identify mutations unique to the cancer. Computational methods predict likely neoantigens and a patient-specific mRNA-based vaccine, intismeran, is manufactured. Highly encouraging results have been observed where the combination of intismeran and anti-PD1 significantly prolong recurrence-free survival in resected melanoma patients [20]. This strategy seems quite adaptable to lung tumors driven by oncogenic RTKs.

## Materials and Methods

### Cell lines and culture

The murine EGFR-mutant cell lines mEGFRdel19.1, mEGFRdel19.2, and mEGFR-L860R.1 were previously described [7]. Novel mEGFR-L860R.2 and mEGFR-L860R.3 cell lines were similarly derived from primary lung tumors induced in mEGFR-L860R GEMMs. All cell lines were cultured in RPMI-1640 growth medium supplemented with 5% fetal bovine serum (Cytiva) and 1% penicillin-streptomycin at 37°C in a humidified 5% CO_2_ incubator. The osimertinib-resistant lines described herein were generated from transplanted lung tumors that were progressing on osimertinib treatment. The tumors were dissected, minced, digested in ACK Lysis Buffer (Gibco) and cultured in RPMI-1640 medium supplemented with 5% FBS and 1% penicillin-streptomycin supplemented with 200nM osimertinib to maintain TKI resistance.

### CRISPR knockout of mC3 in mEGFRdel19.2 and mEGFR-L860R.1 cells

The mC3-positive parental mEGFR cell lines were submitted to CRISPR/Cas9-mediated deletion of mC3 with the crRNA (5’-CCGTCCAGCTCGACCAGGAT-3’) described in Honkanen et al [21]. mEGFRdel19.2 and mEGFR-L860R.1 cells (300,000) were plated in 60mm dishes and cultured for 4hrs. The cerulean targeting crRNA, ATTO550-labelled tracerRNA and recombinant Cas9 were assembled into ribonucleoprotein (RNP) particles according to Integrated DNA Technologies (IDT) manufacturer’s protocol (Iowa, USA). Assembled RNPs were packaged using Lipofectamine RNAiMAX transfection reagent (ThermoFischer Scientific) and the transfection mixture was added to murine EGFR cell lines. After 48-hour, cells were trypsinized, rinsed with 5 mL of PBS, and resuspended into a single-cell suspension in 1mL of PBS. Cells were flow sorted using the MoFlo XDP100 cell sorter for ATTO550 positivity (Beckman Coulter) with the assistance of the University of Colorado Anschutz Flow Cytometry Core. A second flow sort was performed one week later, collecting mC3 negative cells. A third and final flow sort was performed one week later, again collecting mC3 negative cells. For proper compensation of flow cytometry channels, unstained cell lines were used. Once cells were established, visual verification of the lack of mC3-positive cells was performed with a Keyence BZ-X810 fluorescent microscope (Chicago, IL).

### Preparation and culture of lung fibroblasts from mEGFR GEMMs

Lungs from mEGFR GEMMs were perfused with 5 mL of phosphate-buffered saline (PBS)/heparin solution. Both lobes were mechanically dissociated using a razor blade and incubated for 30 minutes at 37°C with 3.2 mg/mL Collagenase Type 2 (Worthington, 43C14117B), 0.75 mg/ml Elastase (Worthington, 33S14652), 0.2 mg/ml Soybean Trypsin Inhibitor (Worthington, S9B11099N), and 40 μg/ml DNAse I (Sigma). The resulting single cell suspensions were filtered through 70μm strainers (BD Biosciences) and washed with FA3 staining buffer (PBS containing 1% FBS, 2mM EDTA, 10mM HEPES). Samples underwent a red blood cell lysis step (0.15 mM NH_4_Cl, 10 mM KHCO_3_, 0.1 mM Na_2_EDTA, pH 7.2), were washed, and filtered through a 40 μm strainer (BD Biosciences). The single cell suspension samples were placed in tissue culture dishes containing RPMI-1640 medium supplemented with 5% FBS and 1% penicillin-streptomycin and cultured for ∼1 week. An adenovirus preparation (4x10^10^ PFU/ml) encoding Cre recombinase driven by the CMV promoter was added to the cell culture medium for 72 hrs and total RNA was purified using Quick-RNA MiniPrep kits (Zymo Research, Irvine, CA).

### Analysis of drug sensitivity in 96-well assays

All murine EGFR cell lines, including parental, mC3 KO and osimertinib-resistant, were plated at 200-600 cells per well in 96-well tissue culture plates and treated in triplicate with TKIs and pharmacological inhibitors at concentrations ranging from 0 to 300 nM. Cell number per well was estimated after 5-7 days of culture, or when the DMSO-control wells reached 80% confluency using Invitrogen’s CyQUANT Direct Cell Proliferation Assay (Invitrogen, Carlsbad, CA) according to manufacturer’s instructions.

### Quantitative RT-PCR methodology

Aliquots (5 μg) of total RNA were reverse transcribed in a volume of 20 μL using Maxima First Strand cDNA Synthesis Kit (Thermo Scientific, Pittsburgh, PA). Aliquots (2 μL) of a 1:5 dilution of the reverse transcription reactions were submitted to quantitative PCR in 10 μL reactions with SYBR Select Master Mix for CFX (Thermo Fisher Scientific) using a CFX Connect Real-Time PCR Detection System (BioRad, Hercules, CA). The following PCR primer pairs were used: mC3 (Fwd: 5’-TGGAGTACAACGCCATCCAC-3’, Rev: 5’-AACTCCAGCAGGACCATGTG-3’), murine MET (Fwd: 5’-GTTGTCCTTGGTGCAGAGGA-3’, Rev: 5’-CGTGAAGTTGGGGAGCTGAT-3’); murine HGF (Fwd, 5’-AGCTACAGAGGTCCCATGGA-3’, Rev, 5’-CAAGAACTTGTGCCGGTGTG-3’); murine GAPDH (Fwd: 5’-CGTGGAGTCTACTGGCGTCTTCAC-3’, Rev: 5’-CGGAGATGATGACCCTTTTGGC-3’). The mRNA levels were normalized to GAPDH mRNA measured in the same samples and presented as “Relative Expression”.

### Orthotopic mouse lung tumor methods

All procedures and manipulations were performed under protocols approved by the Institutional Animal Care and Use Committees at the Rocky Mountain Regional VA Medical Center and University of Colorado Anschutz Medical Campus. Female C57BL/6J (#000664) and Rag1KO (#002216) mice were purchased from Jackson Laboratory (Bar Harbor, ME). The mEGFRdel19 and mEGFR-L860R GEMMs [7] were generated from in-house breeder colonies. GFP-transgenic mice (C57BL/6-Tg(UBC-GFP)30Scha/J) were generated from an in-house colony established from a breeder pair purchased from Jackson Laboratory (#004353). Murine EGFR mutant lung cancer cells were suspended in a solution of 1.35 mg/mL Matrigel (Corning #354234) diluted in Hank’s Balanced Salt Solution (Corning # 21-020-CV) for injection. Mice were anesthetized with isoflurane, the left side of the mouse was shaved, and a 1 mm incision was made to visualize the ribs and left lobe of the lung. Using a 30-gauge needle, 500,000 cells were injected in 40 µL of matrigel cell mix directly into the left lobe of the lung, the incision closed with staples and removed a week later. Tumors were permitted to establish for 10 days before the mice underwent micro-CT (µCT) imaging on a SkyScan 1276 (BRUKER, Billerica, MA) to obtain pre-treatment tumor volumes. Tumor-bearing mice were treated with 5 mg/kg osimertinib by oral gavage for 5 consecutive days followed by 2 days off. To monitor effects of drug treatment on tumor volume, mice underwent weekly µCT imaging and tumor volume was quantified with ITK-SNAP software (www.itksnap.org).

### Pharmacological agents

Osimertinib and crizotinib were generously provided by patient donation through the Thoracic Oncology unit, University of Colorado Hospital. Afatinib, trametinib and RMC-4550 were purchased from MedChemExpress (Monmouth Junction, NJ).

### Statistics

All graphing and statistical analyses were performed using GraphPad Prism version 10.4. Data are presented as the mean ± standard error of the mean (SEM). Dose-response data were curve fitted using Prism 10 nonlinear curve-fitting and the log(inhibitor) vs. normalized response model with robust regression to calculate IC_50_ values.

## List of Abbreviations

RTK: receptor tyrosine kinase
TKI: tyrosine kinase inhibitor
LUAD: lung adenocarcinoma
EGFR: epidermal growth factor receptor
mC3: mCerulean3
GFP: green fluorescent protein
MET: Mesenchymal Epithelial Transition
HGF: hepatocyte growth factor
ALK: anaplastic lymphoma kinase
μCT: microcomputed tomography

## Declarations

Ethics approval and consent to participate: All experiments were approved by the Institutional Animal Care and Use Committees at the University of Colorado Anschutz Medical Campus and the Rocky Mountain Regional VA Medical Center under approved protocols.

## Author Contributions

Concept and experimental design: JTA, TKH, ACN, RAN, LEH Developed methodology: JTA, TKH, KJK, RAN, LEH Performed experiments and acquired data: JTA, TKH, ACN, LEH Analyzed and interpreted data: JTA, TKH, ACN, RAN, LEH Prepared figures and drafted manuscript: JTA, TKH, LEH Edited and revised manuscripts: JTA, TKH, ACN, KJK, RAN, LEH

## Availability of Data

## Acknowledgements

The authors acknowledge the Flow Cytometry shared resource within the University of Colorado Cancer Center.

